# Habitat types and megabenthos composition from three sponge-dominated high-Arctic seamounts

**DOI:** 10.1101/2022.06.26.497630

**Authors:** Tanja Stratmann, Erik Simon-Lledó, Teresa Maria Morganti, Andrey Vedenin, Autun Purser

## Abstract

Seamounts are isolated underwater mountains often stretching >1,000 m above the seafloor. They are usually identified as biodiversity hotspots of marine life, and host benthos assemblages that may vary on regional (among seamounts) and local (within seamounts) scales. Here, we collected seafloor imagery of three seamounts at the Langseth Ridge in the central Arctic Ocean to assess habitats and megabenthos community composition at the Central Mount (CM), the Karasik Seamount (KS), and the Northern Mount (NM). The majority of seafloor across these seamounts comprised bare rock, covered with a mixed layer consisting of sponge spicule mat intermixed with detrital debris composed primarily of polychaete tubes as well as sand, gravel, and/ or rocks. The megabenthos assemblages consisted of in total 15 invertebrate epibenthos taxa and four fish taxa, contributing to mean megabenthos densities of 55,745 ind. ha^-1^ at CM, 110,442 ind. ha^-1^ at KS, and 65,849 ind. ha^-1^ at NM. The faunal assemblages at all three seamounts were dominated by demosponges of the order Tetractinellida that contributed between 66% (KS) and 85% (CM) to all megabenthos. Megabenthos assemblages living on bare rock or on mixed substrate differed among seamounts and across individual seamounts.

## Introduction

Seamounts are isolated subaquatic mountains of (mostly) volcanic origin that rise at least 1,000 m above the surrounding seafloor ^1^. With a global abundance of ∼10,000 ^2^ to ∼125,000 ^3^ seamounts, they cover a minimum ∼8,000,000 km^2^ ^2^ and form one of the largest biomes on our planet ^4^. Seamounts are often hotspots of deep-water biodiversity ^5–8^ and can support higher species abundances than surrounding continental margins and continental slopes ^9^. This phenomenon is known as the ‘seamount oasis hypothesis’ ^9^ that asserts that benthic invertebrates occur in higher densities and biomasses on seamounts than in other habitats in the deep sea ^9^. Seamounts can also influence the overlying water column and affect the microbial community ^10,11^, phytoplankton ^12^, zooplankton ^13^, and ultimately large fish ^1^, a phenomenon known as the ‘seamount effect’ ^13^.

Regional variations in benthic assemblages among seamounts can be driven by differences in latitude ^14–16^, longitude ^17^, food supply ^18^, water depth ^15,16,19–21^, or distance from shore ^22^. Structuring factors can also be region-specific, for instance Boschen et al. ^23^ identified magnetivity (*i.e*., a proxy for hydrothermal activity ^24^) as main driver of differences in the benthic composition among three seamounts in New Zealand. Overall, many factors known to drive biological community variability in the deep sea are related with water depth (*i.e*., temperature, pressure, oxygen concentration, or food-availability), which restricts most benthic fauna to a limited bathymetric range (*e.g*., ^25^). As a result, available habitat for a given population or community can be fragmented across the seamounts (and continental slopes, *e.g*., ^26^) within a region ^27^, particularly in areas with high variability in seamount vertical sizes (*i.e*., water depth at summits). In addition, most deep-sea benthic fauna are thought to exhibit a biphasic life cycle between the release of pelagic planktotrophic or lecithotrophic larvae (*i.e*., respectively plankton-, or yolk-feeding, *e.g*., ^28^) to the water column for dispersal and subsequent settlement on the substrate for benthic/sessile adult stage. As such, connectivity between species populations on different seamounts is thought to be largely controlled by regional factors affecting larvae transport, development, or resilience, such as food availability and temperature within the water column, hydrographic retention mechanisms, or the presence of suitable habitat where propagules ultimately settle ^27^. However, regional gradients and larval dispersal dynamics can be strongly modulated by small-scale processes across seamounts ^23,27,29,30^. Consequently, integration of local to regional observational scales is often essential to accurately assess spatial patterns in seamount communities.

Variations in benthic assemblages within seamounts have been related to changes in seabed composition (*e.g*., hard substrate availability) ^16,19,23,31^, slope ^16,32^, currents ^32,33^, or food supply ^32^; all factors that typically covary with water depth ^19,23,33,34^. As a result, community composition tends to be depth-stratified within seamounts ^6,27^. However, essential niche requirements, such as a minimum rate of food supply ^35^ or the presence of hard substrate ^36,37^ for many sessile taxa, can be strongly modulated by the topographical complexity of a seamount in interaction with the surrounding water masses (*e.g*., ^38^). Similarly, aggregations of framework-building fauna growing in suitable areas within a seamount can generate a habitat for other species, such as is the case with cold-water coral reefs ^35,39^ and sponge grounds ^40,41^, enhancing local habitat heterogeneity rates and thereby increasing alpha diversity rates ^42^. Given this wealth of possible drivers operating locally, and cumulative effects from their interactions, benthic assemblages within a seamount tend to exhibit rapid shifts in composition across space ^43^, usually reflecting the ranges of one or more environmental gradients that define the boundaries of different habitat types.

There are likely more than 300 seamounts beneath Arctic waters ^44^, but only a few have been the subjects of ecological investigations. Sponge grounds appear to be one of the most commonly found seamount habitats in high latitudes (*e.g*., ^45^), as they have also been observed on N Atlantic seamounts (*e.g*., 40° to 75° N latitude belt; ^46,47^). Roberts et al. ^48^ suggested that short-timescale environmental variability combined with the generally nutrient rich sub-surface water masses generated at the interface of intermediate and deep-sea water masses ^49^ might potentially enhance the development of sponge aggregations and other megafauna (*e.g*., ascidians, cnidarians, echinoderms, and demersal fish) at the Schulz Bank seamount (73.5° N, previously called Schultz Massif seamount ^48^) on the Arctic Mid-Ocean Ridge, between the Greenland Sea and the Norwegian Sea. However, very little is known about the diversity of benthic communities in seamounts at higher Arctic latitudes (*e.g*., >75° N; see also ^50^). Hence, less if anything is known about what are the key factors structuring Arctic seamount communities across different spatial scales, *e.g*., do these differ from those shaping lower latitude seamount communities?

In this study, we used seabed imagery to investigate regional (between seamounts) and local (within seamounts) variations in benthic megafauna communities across the Langseth Ridge in the Central Arctic (86.6° − 86.9° N; approximate water depth 600 – 2000 m). This is a chain of three seamounts (Central Mount, Karasik Seamount, and Northern Mount), where surprisingly dense aggregations of mobile sponges (an astonishing unforeseen trait; ^51^) were recently linked with the consumption of underlying fossilized remnants of an extinct seep community by sponge-associated microorganisms ^50^. We assessed whether areas with different seabed composition (Fig. 1, five potential habitats: H1, dense sponge gardens; H2, mats of extensive polychaeta tubes and sponge spicules covered with sulfide precipitates; H3, sediment with gravel; H4, bare rock; and H5, mixed or undominated substrate) harbored distinct megabenthic assemblages, both within and across seamounts. We propose how different drivers (including sponge enhanced biomass) might interact to structure these remote and largely unexplored deep-sea communities.

**Figure 1.**
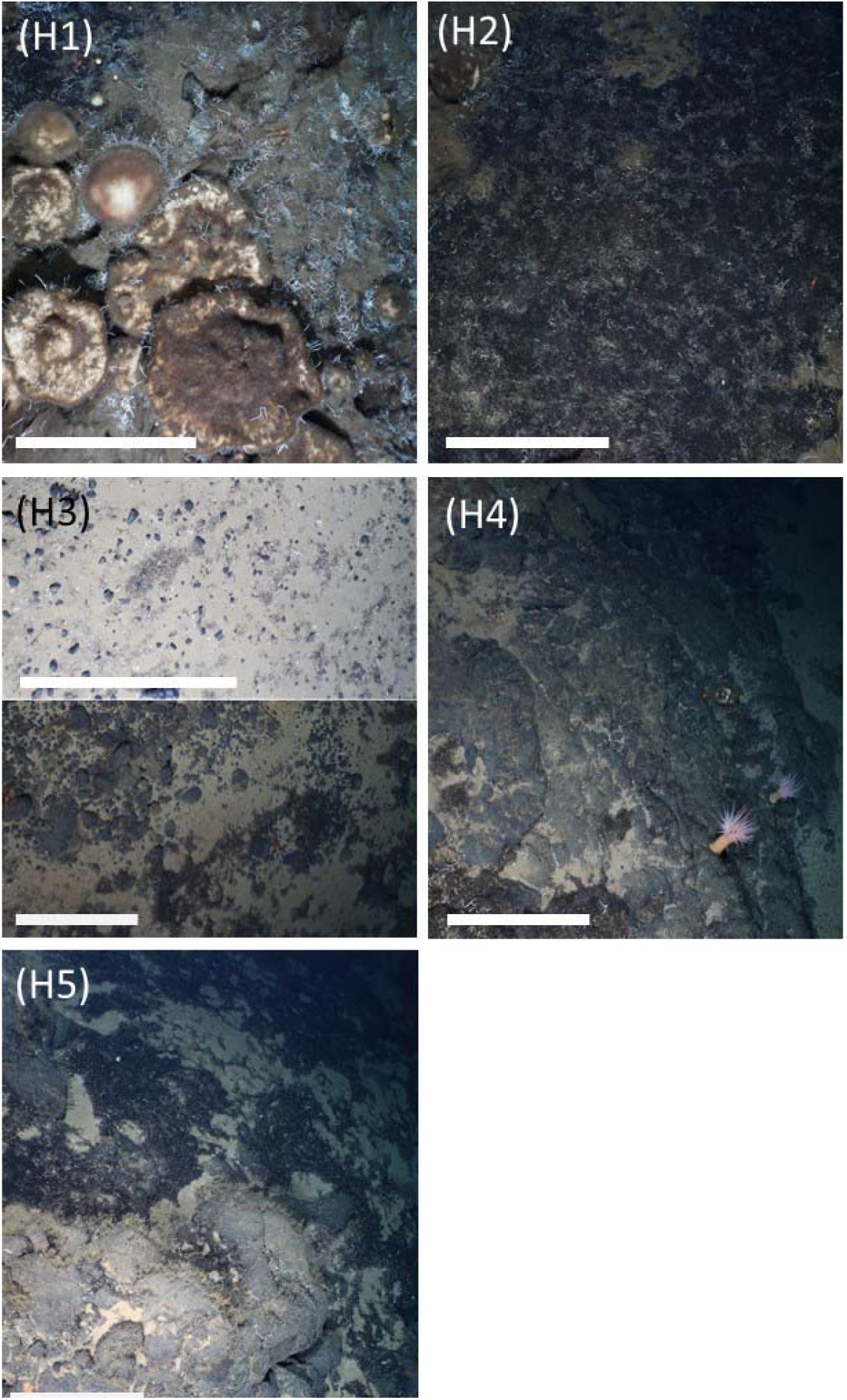
Habitat types identified over the three seamounts: (H1) dense sponge gardens of Tetractinellida gen. indet. sponges extending on top of sponge-produced spicule mats, (H2) mats of Serpulidae indet. and Siboglinidae indet. tubes intermixed with sponge spicules covered with sulfide precipitates, (H3) sediment with gravel, (H4) bare rock, and (H5) mixed substrate. The white bar represents 50 cm.

## Results

### Micro- and macrohabitat types

Most of the seafloor at the Central Mount (CM) was covered with bare rock (habitat type H4, 56% seabed coverage; Table S1, Fig. 1) and mixed substrate (H5, 30.9% seabed coverage; Table S1, Fig. 1), *i.e*., a mixed assemblage of sponge grounds with spicule mats, mats of polychaete tubes, sand with gravel, and/ or bare rocks. Habitat type H2, *i.e*., mats of Serpulidae gen. indet. and Siboglinidae gen. indet. tubes intermixed with sponge spicules and covered with sulfide precipitates, was not observed. The Karasik Seamount (KS) was dominantly covered by H4 (73.4%; Table S1) and the Northern Mount’s (NM) seabed consisted to 56.7% of habitat type H5 and to 30.6% of habitat type H4 (Table S1).

### Quantitative assessment

#### Variations in faunal density

Megabenthos density exhibited substantial variations across the different areas investigated, both at the regional (between seamounts) and at the local (between habitats) scales. Mean faunal density at KS (mean density: 110,442 ind. ha^-1^; 95% confidence intervals: 86,253 − 139,541 ind. ha^-1^) was substantially larger than at CM (mean density: 55,745 ind. ha^-1^; 95% confidence intervals: 43,305 – 71,984 ind. ha^-1^) (Fig. 2A), whereas the assemblages in NM exhibited a large variability, ranging from 34,162 to 119,762 ind. ha^-1^ (in ca. 1,000 specimen samples). Local assessments revealed that the high variability observed at NM was predominantly caused by large differences in faunal density between habitat type H4 (mean density: 113,118 ind. ha^-1^) and H5 (mean density: 14,123 ind. ha^-1^) (Fig. 3A). Densities were consistently smaller in H5 compared to H4 areas, but substantially different densities were also found between habitats of the same type across different seamounts (Fig. 3A), suggesting the existence of faunal density drivers operating at both local and regional scales. Taxon-specific densities specifically for H4 and H5 are presented in Table S3.

**Figure 2.**
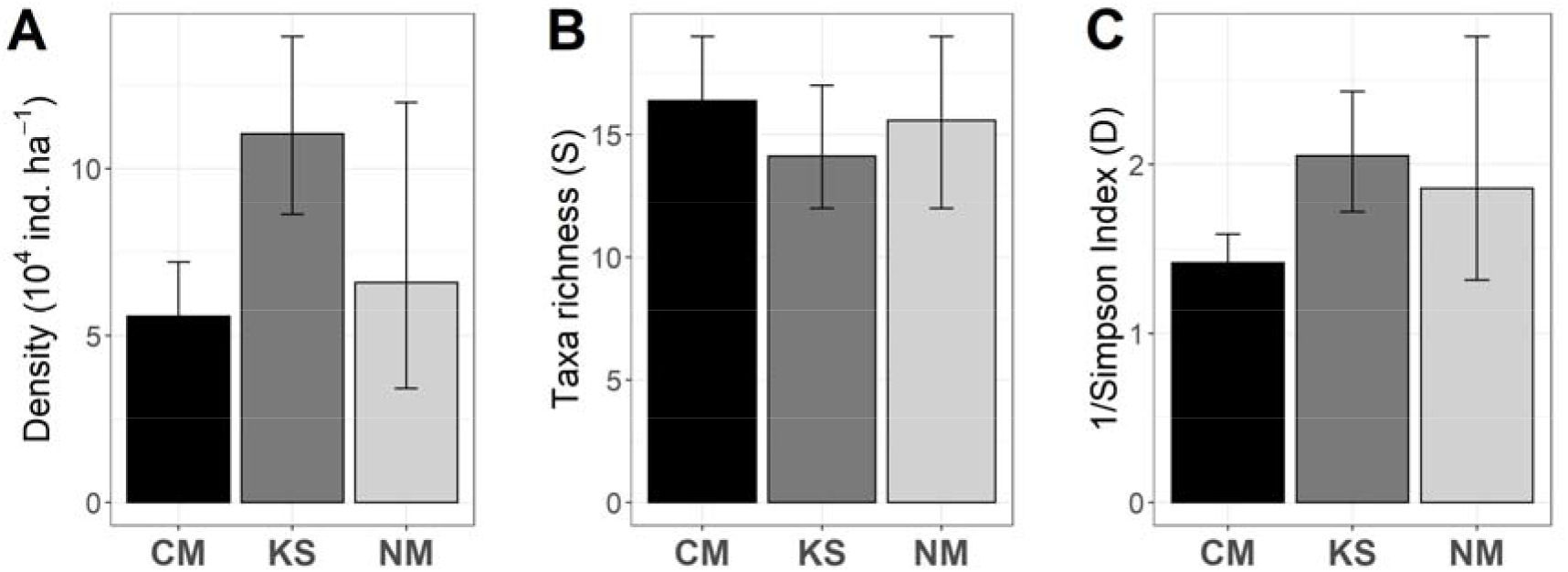
Regional variations in (A) megabenthos density (ind. ha^-1^), (B) taxa richness *S* (in ca. 1,000 specimens), and (C) 1/Simpson index *D* (in ca. 1000 specimens) across the three Arctic seamounts (CM = Central Mount, KS = Karasik Seamount, NM = Northern Mount) investigated. Bars indicate mean values across bootstrap-like sample sets surveyed in each study area. Error bars represent 95% confidence intervals.

**Figure 3.**
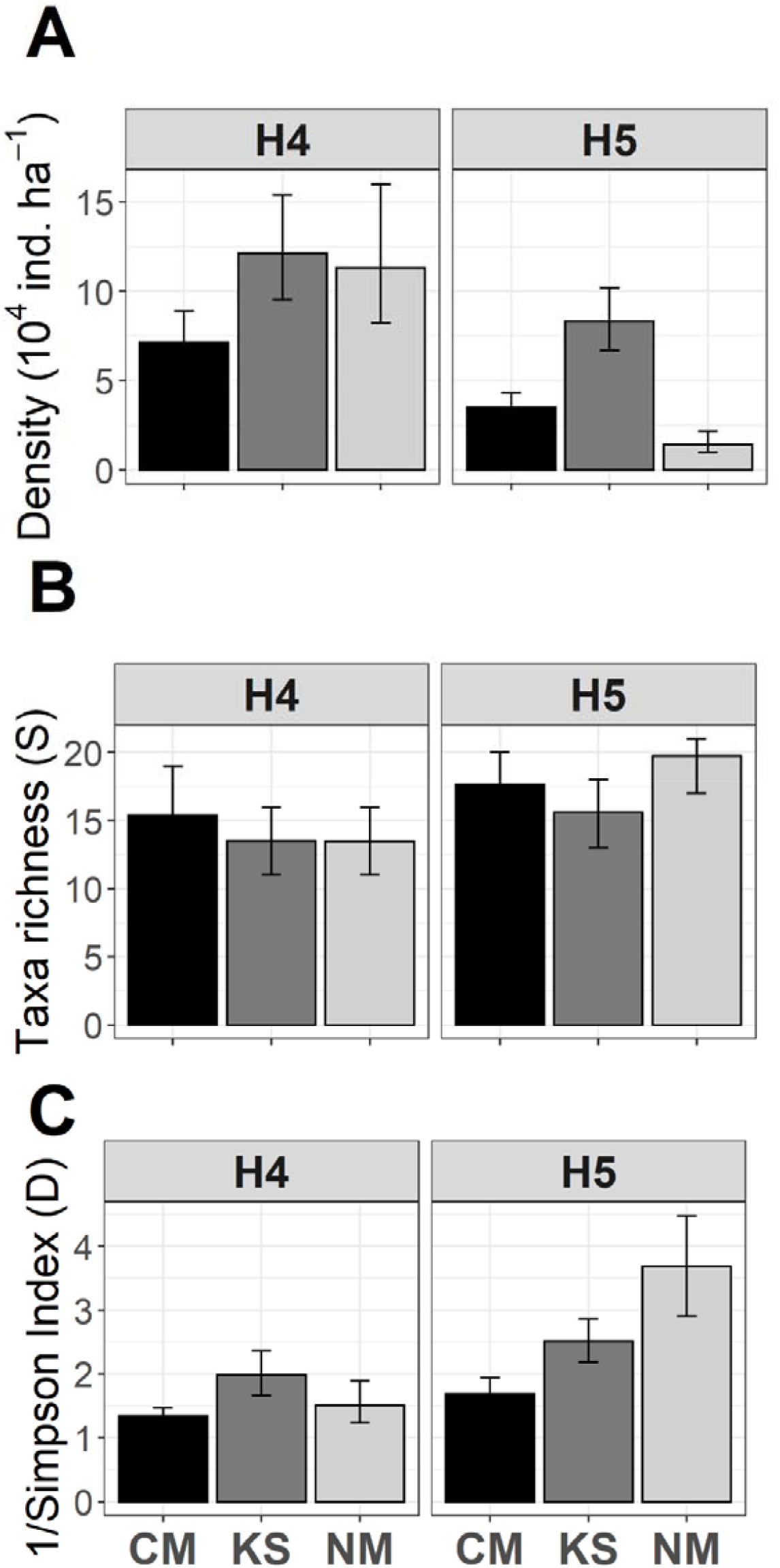
Local variations in (A) megabenthos density (ind. ha^-1^), (B) taxa richness *S* (in ca. 1,000 specimens), and (C) 1/Simpson index *D* (in ca. 1,000 specimens) across the three Arctic Seamounts (CM = Central Mount, KS = Karasik Seamount, NM = Northern Mount) investigated. Bars indicate mean values across bootstrap-like sample sets surveyed in the predominant habitat types (H4 and H5) within each study area. Error bars represent 95% confidence intervals.

#### Variations in diversity

No substantial variations in taxa richness were observed across the different areas investigated, neither regionally nor locally (Fig. 2B, 3B). In contrast, heterogeneity diversity (*i.e*., 1/*D*, an index more sensitive to taxa evenness) was substantially higher at KS than at CM, whereas the assemblages at NM exhibited a large variability for this parameter (Fig. 2C). Similar to faunal density, local assessments revealed that the high variability observed at NM was predominantly caused by large differences in heterogeneity diversity between habitat type H4 (mean 1/*D*: 1.5 effective taxa) and H5 (mean 1/*D*: 3.7 effective taxa) (Fig. 3C). In turn, no major differences were observed in heterogeneity diversity between H4 and H5 areas at CM nor at KS (Fig. 3C). However, heterogeneity diversity in both H4 and H5 areas from KS were consistently higher than in respective H4 and H5 areas from CM, again suggesting the existence of diversity drivers operating at both local and regional scales.

#### Variations in assemblage composition

A total of 15 invertebrate epibenthos taxa and four fish taxa were identified in the image set across the three seamounts studied (Fig. S1). At KS 15 invertebrate and three fish taxa were observed, whereas 14 invertebrates and four fish were detected at CM. At NM all taxa observed on the other seamounts were also found (Table S2).

Multivariate analyses showed substantial variations among the assemblages of different areas investigated, both at the regional (between seamounts) and at the local (between habitats) scales. MDS ordination of regional assemblage composition data readily distinguished the bootstrap-like samples from the three study areas, particularly those from KS and NM (Fig. 4A), as assemblage dissimilarity was higher between these two areas (β_BC_: 32.1%) than between CM and the other two seamounts (β_BC_: 24.6 – 25%). A much larger variation was found, however, within the assemblages of NM than within both CM and KS assemblages (Fig. 4A). Local assessments revealed that the high within-sample variability observed at NM was predominantly caused by a high dissimilarity between the assemblages in H5 and H4 (β_BC_: 31.3%; Fig. 4B), the latter exhibiting a high resemblance with the assemblage from H4 areas at CM (β_BC_: 22.3%). In contrast, dissimilarity between the assemblages of H4 and H5 was less pronounced within CM (β_BC_: 17.3%), and almost inexistent within KS (β_BC_: 13.8%; Fig. 4B, overlapping confidence intervals), suggesting a stronger control of regional drivers at CM and KS areas, and a stronger control of local drivers at NM area.

**Figure 4.**
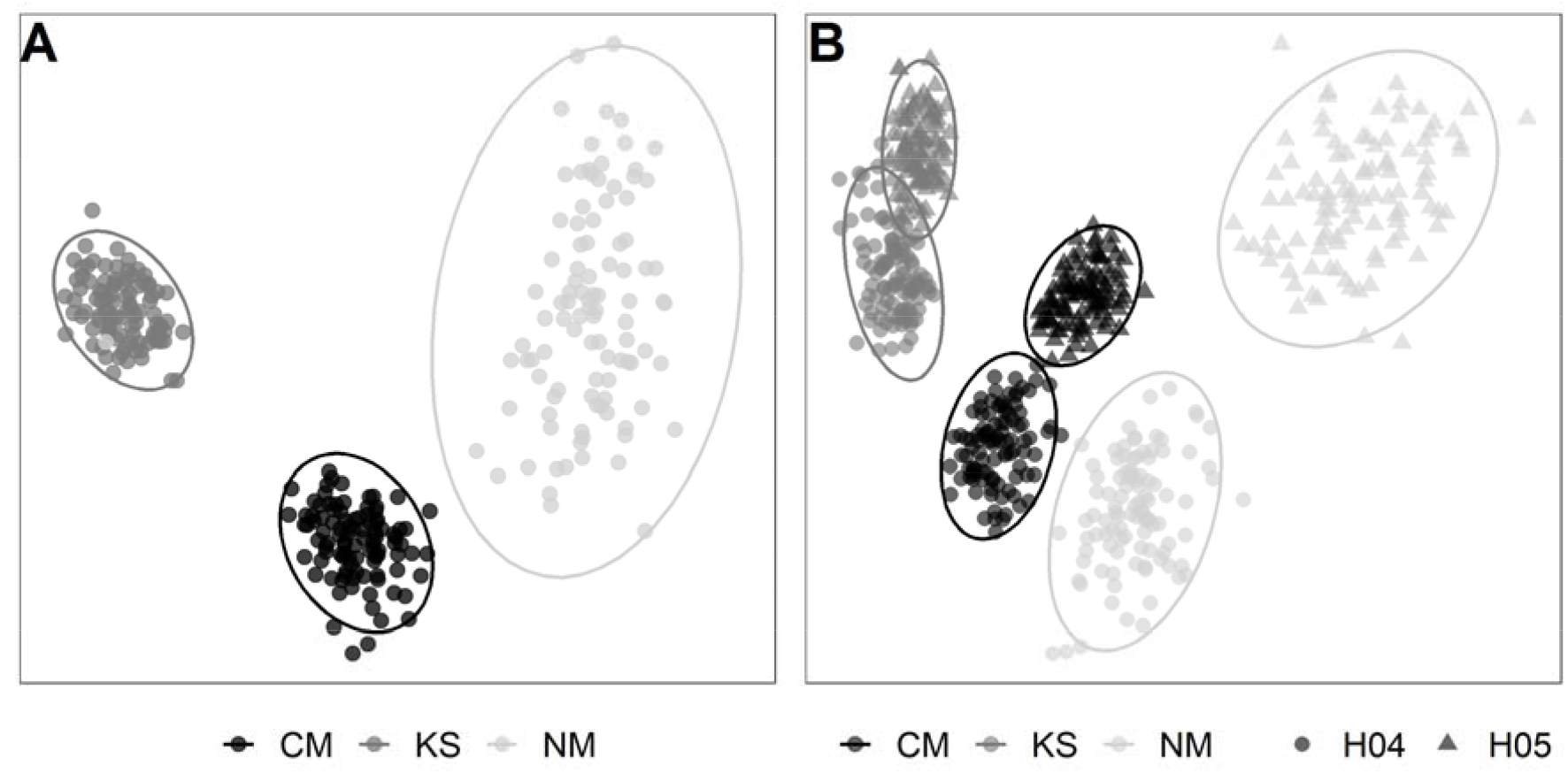
nMDS plots showing regional (between seamounts) and local variations (between habitat types) in faunal assemblage composition. (A) Regional assessment based on 100 randomly selected bootstrap-like samples for each study area (seamounts: CM = Central Mount, KS = Karasik Seamount, and NM = Northern Mount) (nMDS stress: 0.09). (B) Local assessment based on 100 randomly selected bootstrap-like samples for each habitat type (H4 = bare rock, H5 = mixed substrate) in each study area (nMDS stress: 0.12). Ellipses represent 95% confidence intervals.

### Qualitative assessment

#### Faunal assemblage at the Central Mount

The faunal assemblage at CM was clearly dominated by the sponges of the order Tetractinellida gen. indet. (84.9% of all fauna; 23,070 ind. ha^-1^ sponges with <8 cm diameter, 24,348 ind. ha^-1^ sponges with >8 cm diameter; Fig. 5A, Table S2). The second, third, and fourth most abundant taxa were, respectively: the shrimp *Bythocaris* sp. indet. (3,502 ind. ha^-1^; 6.27% of all fauna), the brittle star *Ophiostriatus striatus* sp. inc. (1,133 ind. ha^-1^; 2.03% of all fauna), and the polychaetes *Apomatus globifer* sp. inc./ *Hyalopomatus claparedii* sp. inc. (1,071 ind. ha^-1^; 1.92% of all fauna) (Fig. 5A). All other fauna accounted for 4.83% of the total faunal density.

**Figure 5.**
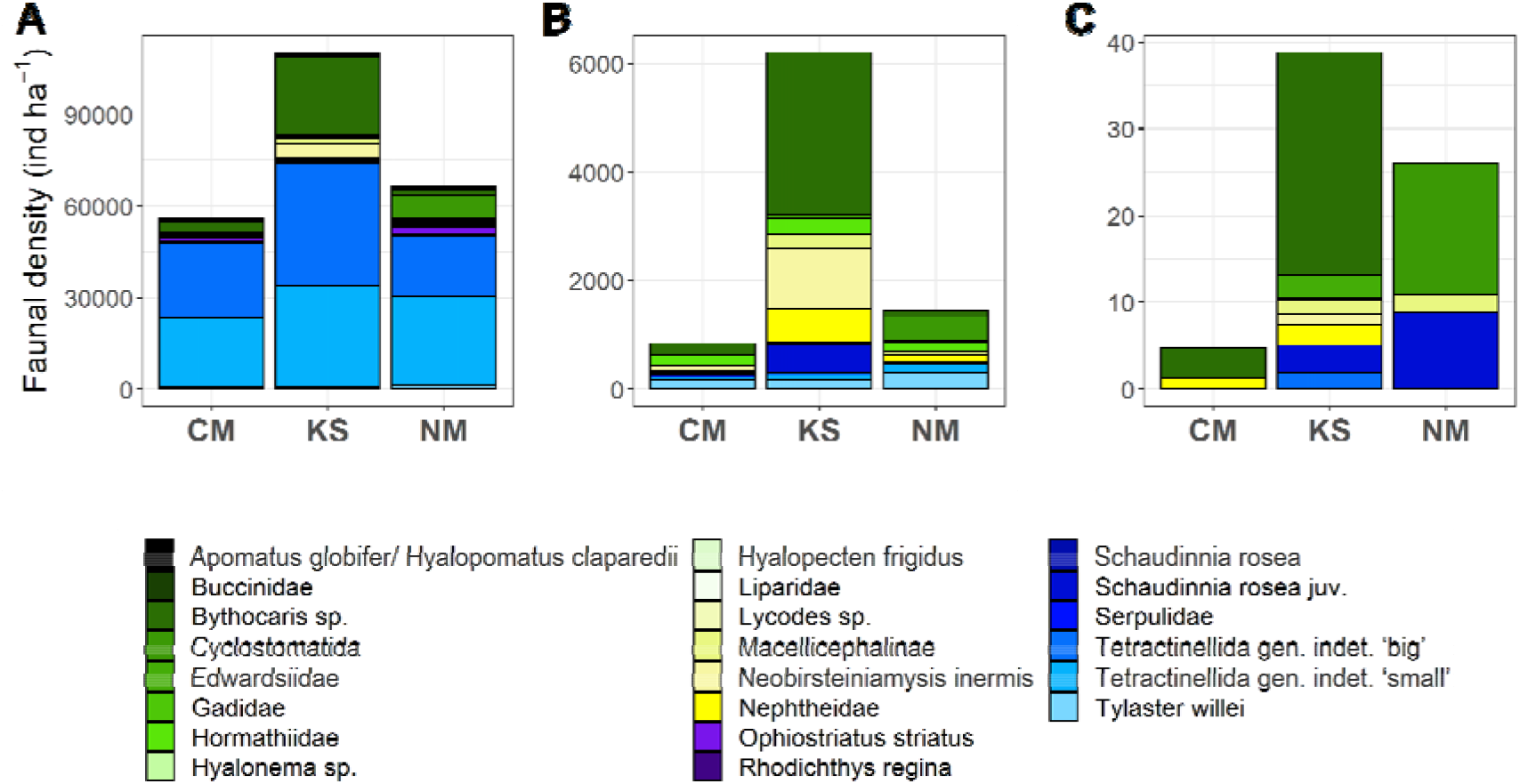
Invertebrate epibenthic megabenthos and fish densities (ind. ha^-1^) at Central Mount (CM), Karasik Seamount (KS), and Northern Mount (NM). (A) All fauna observed at the different seamounts, (B) all fauna that was observed being physically attached to Tetractinellida gen. indet. or walking/ crawling on top of Tetractinellida gen. indet., (C) all fauna that was physically associated with Hexactinellida or was crawling on top of Hexactinellida.

A total of 852 specimens (1.53% of all fauna) were found associated with sponges, *i.e*., either as attached sessile epifauna or as mobile epifauna crawling over these, at CM. Among these, only five specimens were found associated with Hexactinellida sponges, with the remainder found on Tetractinellida specimens. The taxa most frequently associated with Tetractinellida gen. indet. were the shrimp *Bythocaris* sp. indet. (212 ind. ha^-1^; 25.1% of fauna associated with Tetractinellida gen. indet.), the anemone Hormathiidae gen. indet. (185 ind. ha^-1^; 21.8% of fauna associated with Tetractinellida gen. indet.), and the starfish *Tylaster willei* sp. inc. (180 ind. ha^-1^; 21.2% of fauna associated with Tetractinellida gen. indet.) (Fig. 5B-C).

#### Faunal assemblage at the Karasik Seamount

The faunal assemblage at KS was dominated by the sponges of the order Tetractinellida (66.4% of all fauna; 33,422 ind. ha^-1^ sponges with <8 cm diameter, 39,860 ind. ha^-1^ sponges with >8 cm diameter; Fig. 5A) and the shrimp *Bythocaris* sp. indet. (22.9% of all fauna; 25,284 ind. ha^-1^; Fig. 5A, Table S2). The remaining faunal assemblage was predominantly composed by the mysid *Neobirsteiniamysis inermis* sp. inc. (4,360 ind. ha^-1^), and the polychaete Macellicephalinae gen. indet. (2,054 ind. ha^-1^) and *Apomatus globifer* sp. inc./ *Hyalopomatus claparedii* sp. inc. (1,391 ind. ha^-1^). All other fauna accounted for 2.86% of the total faunal density.

A total of 6,244 specimens (5.66% of all fauna) were found associated with sponges, *i.e*., either attached, crawling over or feeding on these, at KS. Among these, only 39 specimens were found associated to Hexactinellida sponges and the rest were on Tetractinellida sponges. The taxa most frequently associated with Tetractinellida gen. indet. were the shrimp *Bythocaris* sp. indet. (2,975 ind. ha^-1^; 47.9% of fauna associated with Tetractinellida gen. indet.), the mysid *Neobirsteiniamysis inermis* sp. inc. (1,129 ind. ha^-1^; 18.2% of fauna associated with Tetractinellida gen. indet.), the soft coral Nephtheidae gen. indet. (623 ind. ha^-1^; 10.0% of fauna associated with Tetractinellida gen. indet.), and juveniles of the sponge *Schaudinnia rosea* sp. inc. (527 ind. ha^-1^; 8.49% of fauna associated with Tetractinellida gen. indet.) (Fig.5B-C).

#### Faunal assemblage at the Northern Mount

The faunal assemblage at NM was dominated by the sponge of the order Tetractinellida gen. indet. (73.9% of all fauna; 28,825 ind. ha^-1^ sponges with <8 cm diameter, 20,054 ind. ha^-1^ sponges with >8 cm diameter; Fig. 5A; Table S2) and the bryzoan Cyclostomatida fam. indet. (7,469 ind. ha^-1^; 11.3% of all fauna). The remaining faunal assemblage was predominantly composed by the brittle star *Ophiostriatus striatus* sp. inc. (2,380 ind. ha^-1^), the shrimp *Bythocaris* sp. indet. (2,272 ind. ha^-1^), the starfish *Tylaster willei* sp. inc. (1,435 ind. ha^-1^), and the polychaete Macellicephalinae gen. indet. (973 ind. ha^-1^). All other fauna accounted for 4.01% of the total faunal density.

A total of 1,484 specimens (2.25% of all fauna) were found associated with sponges, *i.e*., either attached, crawling over or feeding on these, at NM. Among these, only 26 specimens were found associated to Hexactinellida sponges with the rest on Tetractinellida gen. indet. sponges. The taxa most frequently associated with Tetractinellida gen. indet., were the bryozoan Cyclostomatida fam. indet. (445 ind. ha^-1^, 30.5% of fauna associated with Tetractinellida gen. indet.), the starfish *Tylaster willei* sp. inc. (303 ind. ha^-1^, 20.8%), other small Tetractinellida gen. indet. specimens (167 ind. ha^-1^, 11.4%), and anemone Hormathiidae gen. indet. (161 ind. ha^-1^, 11.1%) (Fig. 5B-C).

## Discussion

The seafloor at the three seamounts of the Langseth Ridge consisted mainly of bare rock, sand, and gravel along with a mix of biogenic structures composed of reef-forming sponge grounds, spicule mats, and polychaete tubes. Our results showed that the megafaunal densities and assemblage composition, but not taxon richness, differed at regional (between seamounts) and local (within seamount, between habitats) scales across the studied area. Demosponges of the order Tetractinellida numerically dominated the assemblages across the three seamounts, possibly owing to their unique capacity to source carbon directly from the refractory matter available on the seabed ^50^, which likely makes these mostly (if not fully) independent from the water column food-supply in such a low primary productivity area ^52,53^. Shrimps (*Bythocaris* sp. indet.; Central Mount, Karasik Seamount) and bryozoans (Cyclostomatida fam. indet.; Northern Mount) were the other most abundant taxa encountered, yet at far from the densities exhibited by Tetractinellida sponges in a much smaller area. Here, we discuss the potential processes causing the observed variations in megabenthic composition at different scales and the role that Tetractinellida sponges may play in these high Arctic sponge grounds.

Variations in assemblage composition observed in regional assessments were likely related to inherent differences in megabenthos density across the three seamounts; the highest density was observed at the Karasik Seamount and the lowest at the Central Mount. This difference in densities could be correlated with the height of the seamounts: the Karasik Seamount is the highest seamount (summit depth: 585 m), followed by the Northern Mount (summit depth: 631 m), and the Central Mount (summit depth: 722 m). Depth, or more precisely, the strong covariation of key factors (*i.e*., food supply and temperature) with increasing depth ^54^ has been widely highlighted as major proxy for deep-sea benthic abundance and biomass ^55,56^. As such, and in line with our results, many studies have shown how depth-related variations in population densities can yield markedly distinct benthic communities in seamounts ^18,22,57^.

Water temperature and current strength may be other drivers of the variations in faunal abundance observed. For instance, water temperatures measured at the Karasik Seamount (0.66°C) and the Northern Mount (0.68°C) were higher than at the Central Mount (0.23°C) ^58^. In contrast, current velocity measured during the cruise were generally weak (<0.1 cm s^-1^) with a predominantly westwards component and no evidence of associated upwelling currents ^59,60^. It is hence more plausible that food supply and temperature decreases with depth have a stronger influence on the observed variations in megabenthic abundance than the overlying current dynamics. It is noted that bottom currents and hydrographic processes can typically exhibit periodic or seasonal increases, leading to enhanced food supply rates (*e.g*., ^61^), particularly in interaction with the complex topography of seamounts ^38^. However, we rule out the possibility that high densities of the bryzoan Cyclostomatida fam. indet. at the Northern Mount was related to increased seasonal currents due to the sluggishness of the current over the year, and the year round ice cover. Variations in assemblage composition at local scale appears to be clearly driven by the existence of different habitats. Habitat type H4 (bare rock) and H5 (mixed substrate composed by sponge spicule mats intermixed with detrital (in)organic polychaete-tube debris, gravel and rocks) covered between 87% and 89% of the seamount areas studied. Habitat type H5 supported 14,123 ind. ha^-1^ (Northern Mount) to 83,082 ind. ha^-1^ (Karasik Seamount), whereas habitat type H4 supported a relatively denser community of 71,362 ind. ha^-1^ (Central Mount) to 121,442 ind. ha^-1^ (Karasik Seamount). This difference in megabenthic densities between H4 and H5 was partly related to variations in morphotype composition between these habitats at the Northern Mount and the Central Mount. For instance, in both seamounts, only very few *Ophiostriatus striatus* sp. inc. brittle stars were observed across bare rocks (H4), whereas their densities ranged from 1,904 ind. ha^-1^ (Central Mount) to 3,888 ind. ha^-1^ (Northern Mount) across the mixed substrate seafloor areas (H5). *O. striatus* is an opportunistic deposit feeder that was observed grazing upon fresh and detrital ice algae in the Nansen Basin close to the Gakkel Ridge during the minimum sea ice extent in 2012 ^62,63^. The mixed substrate, which can be of up to 15 cm thickness at the seamounts, may trap settled particles ^50^ which could subsequently serve as a food source for *O. striatus*. This would explain the very low densities of this brittle star across bare rocks where potentially increased hydrodynamics might prevent detritus accumulation. However, it remains unclear why *O. striatus* is almost absent from the Karasik Seamount as Zhulay and colleagues observed uncommon swimming behavior in this species ^64^, which might facilitate them to swim from the Central Mount to the Karasik Seamount. This suggests that a combination of regional and local environmental differences likely causes the variability between the megabenthos assemblages at the Northern and Central Mount and that at the Karasik Seamount.

Besides the habitats H4 and H5, also habitats H1 (Tetractinellida sponge grounds) and H3 (sediment with gravel) were observed at all seamounts, whereas habitat H2 (mats of polychaete tubes) was found only at the Northern Mount and the Central Mount. However, owing to the primarily exploratory nature of the research cruise to investigate the geological, geochemical, and biological processes of the active hydrothermal vent at the Gakkel Ridge ^65^ and seamounts at the Langseth Ridge in 2016 ^50,51,65^, the surrounding topography and community structure were not well known before the expedition. Therefore, no previous information was available to delineate different environmental strata in order to carry out a balanced sampling effort. Our bootstrap-based assessment allowed the reduction of this study limitation (*i.e*., unbalanced sampling effort across different habitat types) by focusing on the variability associated to different ecological estimators rather than in the actual estimations (*e.g*., mean values), which can be a robust way for instance, in detecting patterns in opportunistic deep-sea datasets (*e.g*., ^66,67^), yet was only conceived here as a preliminary approach. In this regard, the comparably smaller image sample size for habitats H1, H2, and H3 did not allow for reliable statistical comparison of these with the more dominant H4 and H5. Thus, based on our preliminary work, future studies aimed at acquiring a better understand the ecology of this area shall now be able to appropriately design benthic image surveys, *i.e*., yielding even sampling effort across each of the, now characterized, Langseth Ridge habitat types.

Sponge grounds, known as areas where sponges reach densities of 0.5–1 m^-2^ (still image surveys ^68^) to 0.03–0.1 m^-2^ (video surveys ^40^), usually exhibit an increased associated benthic diversity and biomass when compared to adjacent non-sponge habitats ^41^. Sponges typically enhance the complexity of habitats by increasing the (three-dimensional) hard surface area available for other fauna to interact (*e.g*., settle, reproduce, and/or hide ^37,69–72^). For instance, at the Schulz Bank seamount large demosponges like *Geodia* sp. and *Stelletta* sp. typically have ascidians or other sponges (such as the encrusting sponge *Hexadella dedritifera*) growing along their edges ^45^. Similarly, Tetractinellida sponges at the Langseth Ridge host diverse species, such as juveniles of the sponge *Schaudinnia rosea* sp. inc., the anthozoan Hormathiidae gen. indet., the actinarian Edwardsiidae gen. indet., the byrozoan Cyclostomatida fam. indet., the soft coral Nephtheidae gen. indet., or the polychaete *Apomatus globifer* sp. inc./ *Hyalopomatus claparedii* sp. inc. The latter colonized the edge of the Tetractinellida gen. indet. sponges as they may benefit from the water fluxes generated by the pumping activity of the sponges as well as the particle detritus expelled by them as a source of food for the epi-endobiota ^73^. In comparison, only a few specimens were found associated to the hexactinellid sponges at the seamounts of the Langseth Ridge: few sponge specimens belonging to *Schaudinnia rosea* sp. inc. and Tetractinellida gen. indet. sponge specimens, the bryozoan Cyclostomatida fam. indet., the actinarian Edwardsiidae gen. indet., and the soft coral Nephtheidae gen. indet. The difference in the number of associated specimens between the Hexactinellida and Tetractinellida sponges can be related to their different morphology (papillate/globular and massive, respectively) and the spicule “fur” produced by Tetractinellida that facilitate the epifauna settlement ^74^. Additionally, Tetractinellida sponges at the Northern Mount, Karasik Seamount, or Central Mount had a large variety of mobile fauna associated with them, such as the starfish *Tylaster willei* sp. inc., the shrimp *Bythocaris* sp. indet., and the mysid *Neobirsteiniamysis inermis* sp. inc. However, having only seafloor imagery available, we were unable to define whether these taxa predated upon the sponges, used them as three-dimensional structures on which to sit higher in the water column ^42^, crawled over them, or fed on sponge detritus particles ^73^. Such a unique aggregation of sponges (*i.e*., independent from typically limiting water food supply) are likely to drive interesting or even unexpected patterns in the wider community, hence further ecological research at the Langseth Ridge is expected.

In conclusion, this study presents the first detailed description of megabenthos assemblages at the northernmost seamounts explored so far. Interestingly, taxa richness did not differ between seamounts and habitats. While the megabenthos community composition showed substantial differences at regional and local scale, likely driven by intrinsic seamount characteristics (water temperature and depth) and distinct habitats respectively. The Northern Mount had the highest density of bryozoans, which were almost absent in other seamounts and a more pronounced difference in megabenthic composition between the bare rock and mixed substrate habitats. So far, there is no evidence of particular processes, such as the increase of bottom currents or the different hydrographic conditions at the Northern Mount for explaining such distinctive features when compared to the other two seamounts. Further video and/ or image transects at the individual seamounts are required to assess the megabenthos communities inhabiting the three other classified habitats, and to address the possible interactions between mobile fauna and the sponges.

## Materials and Methods

### Study area

The Langseth Ridge is an underwater ice-covered mountain ridge in the central Arctic Ocean that stretches approximately 125 km from 87°N, 62°E to 85°55’N, 57.45’E ^65,75^ (Fig. 6). It is comprised of three summits, the Central Mount, the Karasik Seamount, and the Northern Mount. The Central Mount has its summit at 86°47.83’N, 61°54.52’E where its maximum elevation reaches to 722 m below the sea surface ^65^. This seamount has a gradually increasing slope from 3,300 m to its point of maximum elevation ^65^. Its slope on the western side is steeper than the one on the eastern side where the slope drops to 4,500 m water depth ^65^. The tallest mountain on the Langseth Ridge is the Karasik Seamount which summit is located at 86°43.0’N, 61°17.6’E and reaches to 2,500 m above the seafloor (*i.e*., 585 m below the sea surface ^65^) ^75^. The Northern Mount is located at 86°51.86’N, 61°34’E and has a maximum elevation of 631 m below the sea surface ^65^. This seamount has a steep slope from its peak towards the Gakkel Ridge rift valley in the north at 4,000 m depth ^65^.

**Figure 6.**
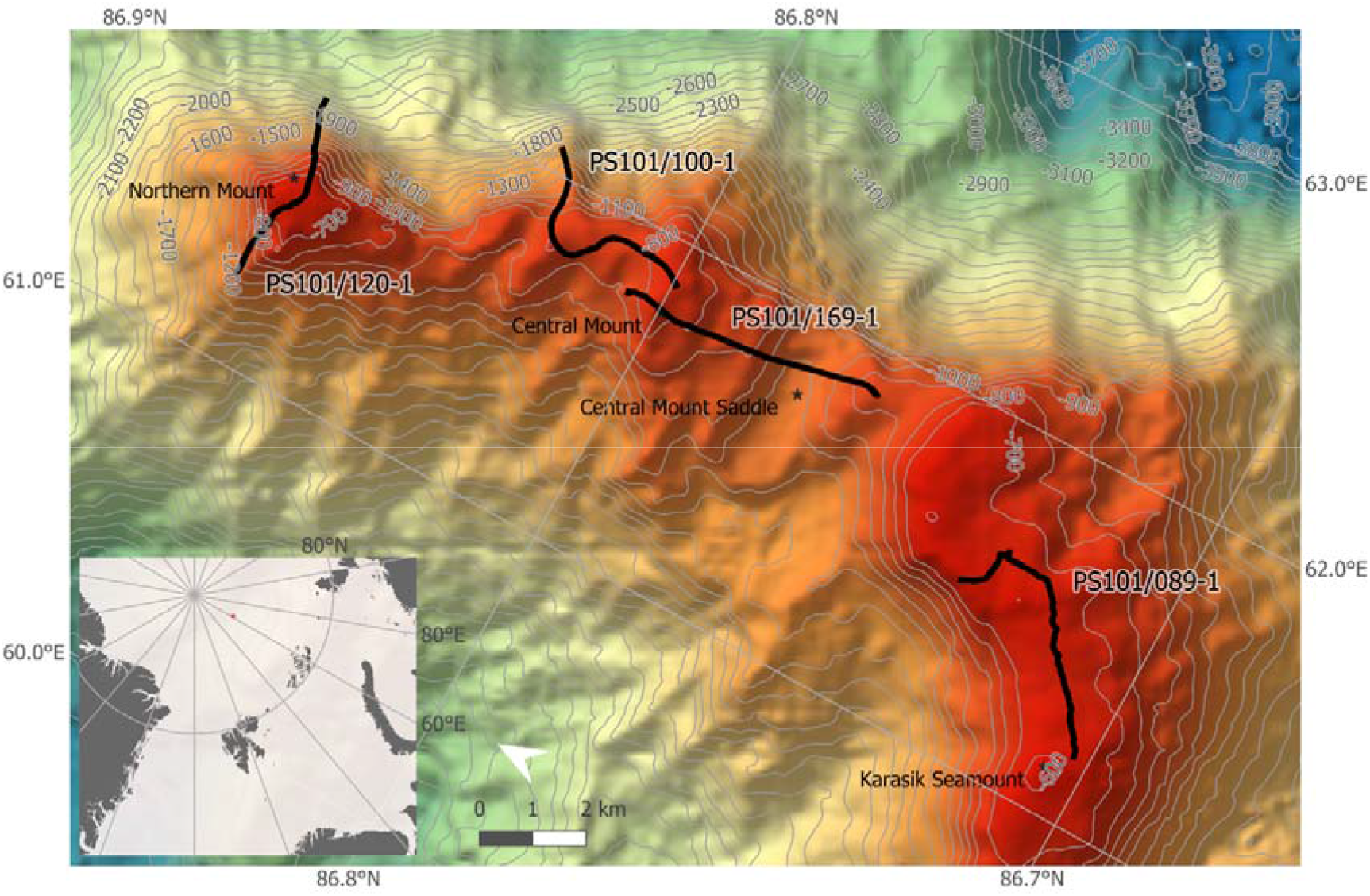
Map of the Northern Mount, Central Mount, and Karasik Seamount along the Langseth Ridge with all Ocean Floor Observation and Bathymetry System (OFOBS) deployments. The station numbers correspond to the OFOBS transect numbers in Table S4.

Bottom water at the three seamounts had a temperature range between -0.02°C (Central Mount) and 0.22°C (Northern Mount) and a salinity of 34.9 PSU ^65^. Oxygen concentration in the same water was only measured at the Northern Mount and amounted to 322 µmol L^-1 76^. Nitrite was not detected in bottom water and ammonium was only found in Karasik Seamount’s bottom water (0.02 µmol L^-1^) ^76^. Phosphate concentrations ranged in bottom water from 0.67 to 0.68 µmol L^-1^, nitrate concentration was between 12.0 and 12.7 µmol L^-1^, and silicate ranged from 5.53 to 5.86 µmol L^-1 76^.

### Seabed image collection and processing

#### Image collection

The high-resolution digital photo camera (CANON EOS 5D Mark III, modified by iSiTEC for underwater applications) of the towed Ocean Floor Observation and Bathymetry System (OFOBS) ^77^ was used to take still images of the seafloor of the three different seamounts. OFOBS was deployed four times (= four transects; Fig. 6 and Table S4) during the RV *Polarstern* cruise PS101 in the central Arctic Ocean (chief scientist: Prof. Dr. Antje Boetius) ^65^. During each deployment, OFOBS was towed 1.5 to 2.5 m above the seafloor at a speed of <1 knot and photographs were taken every 20 seconds to avoid overlap between images. The area of each image was calculated using three laser points on the seafloor that were organized in an equilateral triangle (distance between points: 0.5 m) as reference for scaling (area per image: mean±SE: 8.43±0.17 m^2^). A total of 3,162 photographs were collected, from which 2,099 (17,691 m^2^ of seabed) were used (Table S4), as only bright images collected within the aimed altitude range were selected for analysis. All photographs were loaded into the open-source software “Program for Annotation of Photographs and Rapid Analysis (of Zillions and Zillions) of Images” PAPARA(ZZ)I ^78^.

#### Habitat classification

To classify macro- and microhabitat ^79^ types at the different seamounts, it was recorded for each image whether ≥75% of the seafloor was covered by dense sponge gardens of *Geodia* sp. indet./ *Stelletta* sp. indet. and sponge spicule mats (habitat type H1), by mats of Serpulidae indet. and Siboglinidae indet. tubes covered with sulfide precipitates (habitat type H2), by sediment with gravel (habitat type H3), or by bare rock (habitat type H4). When the seafloor was covered to ≤75% by one of the four habitat types and therefore consisted of an assemblage of sponge ground and spicule mats, mats of polychaete tubes, sediment with gravel, and/ or bare rock, it was classified as mixed substrate (habitat type H5). The five habitat types described in this study are shown in Fig. 1.

#### Biological analysis

Megabenthic fauna (>1 cm size; Fig. S1 visible on the photographs were annotated and identified to the lowest taxonomic hierarchy possible (morphotype [mtp]: typically Genus or Family level). The taxonomic nomenclature of the morphotypes presented follows ^80^.The life-habit of specimens was recorded whenever these were found attached (sessile fauna) or crawling (mobile fauna) on other specimens (*i.e*., generally sponges). As it is not possible to distinguish between *Geodia* sp. indet. and *Stelletta* sp. indet. sponges on seabed images, these specimens were identified as Tetractinellida gen. indet. (Fig. S1H). The latter were annotated based on their diameter size as “Tetractinellida gen. indet. ‘big’” (diameter >8cm) and “Tetractinellida gen. indet. ‘small’” (diameter <8cm). Polychaetes of the family Serpulidae indet. and Siboglinidae indet. (Fig. S1M) were not annotated individually, but as patches. The polychaetes *Apomatus globifer* sp. inc/ *Hyalopomatus claparedii* sp. inc (Fig. S1N) that were observed associated with Tetractinellida gen. indet. specimens were annotated as present/ absent, *i.e*., they were annotated once per image when they were present. The uncertainty of the image-based identifications was indicated following the recommendations by ^80^ for standardization of the open taxonomic nomenclature.

#### Quantitative data analysis

Patterns in diversity and distribution of faunal assemblages were quantitatively assessed at two scales: i) regionally (scale: 10s km), across the three different seamounts (Central Mount, Karasik Seamount, and Northern Mount), and ii) locally (scale: 100s m), between the two habitats with the largest seafloor coverage (H4 and H5; encompassing 94% of all specimens and 88% of all the seabed area surveyed) across the three seamounts. In each case, megabenthos specimen data were pooled for each study area or target stratum, *e.g*., per seamount or habitat type, and then resampled using a modified form of bootstrapping ^81^.

Resampling methods provide robust estimates of variability and confidence intervals of sample parameters ^82,83^, and are particularly well suited to analyze seabed image data obtained from survey designs that lack true sample replication (see *e.g*., ^66,67^). To implement the bootstrap, image data were randomly resampled with replacement until a minimum of 1,000 megabenthos specimens were encompassed, and that process was repeated 10,000 times for each target stratum. This process yielded bootstrap-like samples (bootstrap generated sub-samples) with fixed specimen count size, ranging in total seabed cover from 72 to 490 m^2^, to minimize the potential effect of variable faunal densities in the estimation of ecological parameters.

A range of ecological parameters were calculated from each set of bootstrap-like samples to compare the assemblages from different target strata. Patterns in abundance were assessed by estimation of numerical density (ind. ha^-1^), whereas diversity was assessed by estimation of taxa richness (*S*, in ca. 1,000 specimens) and Simpson’s index (*D*, in ca. 1,000 specimens) ^84^. Variations in assemblage composition were assessed by non-metric multidimensional scaling (nMDS) ordination of bootstrap-like samples, based on the Bray-Curtis dissimilarity (or beta-diversity, β_BC_) measure ^85^ calculated using square-root-transformed faunal density. Mean values of each parameter in each target stratum were calculated, along with corresponding 95% confidence intervals based on the simple percentile method ^81^. All analyses were performed using a custom *R* ^86^ script using multiple functions of the *vegan* package ^87^. Variations in ecological parameters between study areas were reported by comparing 95% confidence intervals (*i.e*., the upper limit of a given estimate must be lower than the lower limits of the estimate that is compared to). Such cases are significant at p < 0.05, but the true (undetermined) p-value will, necessarily, be considerably lower.

## Acknowledgements

We thank chief scientist and project leader Prof. Dr. Antje Boetius (AWI), captain and crew of RV *Polarstern* for their excellent support during research cruise PS101. Laura Hehemann (AWI) is acknowledged for preparing the map of the study area and we are highly grateful for the help of citizen scientist Annette Stratmann with annotating putative seafloor habitats. The research cruise received funding from the DFG Cluster of Excellence “The Ocean in the Earth System at Bremen University (grant no. 49926684) and from the ERC Advanced Grant ABYSS (grant no. 294757) to the chief scientist. TS was supported by the Dutch Research Council NWO (NWO-Rubicon grant no. 019.182EN.012, NWO-Talent program Veni grant no. VI.Veni.212.211), and the publication of this study was enabled by NWO-ALW grant 856.18.003.

## Author contributions

TS conceived the idea, annotated the seafloor images, analyzed the data, and wrote the manuscript. ESL performed the statistical data analysis and wrote the manuscript. TM conceived the idea, revised and edited the manuscript. AV identified the megabenthos species on the seafloor images, revised and edited the manuscript. AP conceived the idea, collected the seafloor images, revised and edited the manuscript.

## Data Availability Statement

All “Ocean Floor Observation and Bathymetry System” (OFOBS) images collected during the RV *Polarstern* PS101 cruise are available at https://doi.pangaea.de/10.1594/PANGAEA.871550. All data generated and analyzed during this study are included in its supplementary information files.

